# Robust and Memoryless Median Estimation for Real-Time Spike Detection

**DOI:** 10.1101/2024.07.19.604279

**Authors:** Ariel Burman, Jordi Solé-Casals, Sergio E. Lew

**Affiliations:** Universidad de Buenos Aires. Instituto de Ingeniería Biomédica. Buenos Aries, Argentina; Data and Signal Processing Group, University of Vic-Central University of Catalonia, 08500 Vic, Spain

## Abstract

We propose a novel moving median estimator specifically designed for online detection of threshold crossings in multi-channel signals, such as extracellular neural recordings. This estimator offers two key advantages: a reduced sensitivity to outliers and the elimination of memory requirements for storing arrival times. Furthermore, its design facilitates parallel implementation on FPGAs, making it ideal for real-time processing of multi-channel recordings.

## Introduction

Median filtering is a widely used technique to remove or detect infrequent events in signal and image processing. Depending on what part of the raw signal is removed, the median filter can be employed to eliminate or to capture impulse noise from signals and images [1, 2], as in the case of extracellular neuronal spikes or “pepper and salt “ noise, respectively.

When those few frequent events are contaminated with Gaussian noise, as in the case of extracellular neuron recordings, an optimal threshold can be calculated to separate spikes from noise [3]. Indeed, most of the state-of-the art neuronal decoding techniques starts with threshold-based spikes detection, independently of what kind of analysis is done after this stage, i.e. multiunit or single unit spike sorting isolation decoding [4–6].

While it is straightforward to compute the median-based threshold in offline spike detection analysis, limitations related to speed and memory appear in real time neuronal decoding of multichannel recordings, mainly due to the huge amount of samples per seconds that arrive from the recording equipment and the non-stationary characteristics of those signals. In consequence, to have a fast and accurate real-time median-driven threshold estimator becomes critical.

In digital systems, sample median computations relies in sorting methods, which are a special case of selection algorithms. In an ordered odd buffer, the buffer middle sample results in the median of those samples. Thus, improving the efficiency of the sample ordering process impacts directly in the estimation speed, *O*(*N* log *N*)) [7].

In a moving (sliding window) median estimator, as soon as the window moves over a new sample, two alternatives can be used in order to estimate the new buffer median value, a naive one that implies to re-order the whole buffer as if it were a new and independent set of samples, or a more efficient one that results from dropping out the oldest sample in the buffer and inserting the new one in such a position that keep the buffer ordered. While the former affects the speed of median estimation, the latter need to keep in memory the temporal order in which every sample arrived to the buffer, requiring as many memory as positions in the buffer exist [2].

We present here an alternative method to compute the median in sliding windows and, in consequence, the optimal moving threshold to detect impulse noise from Gaussian one. The method avoids keeping in memory the order the sample arrival without re-ordering the whole buffer every time a sample comes in. We demonstrate that the proposed method is unbiased and robust compared to the traditional moving median, particularly when the underlying process is stationary. Furthermore, it adapts to a new sample distribution in at most 1.5 times the time required by the traditional method during significant shifts in the signal baseline, such as those encountered in extracellular recordings due to electrode movements.

## Materials and methods

Let *X*(*t*) be a continuous random process, with probability distribution density function *f*_*X*_ (*x*), which we will only assume it is uni-modal. The classical moving median estimator can be computed, at time *t*_*n*_ as

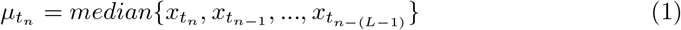

when using a buffer of length *L*.

A naive implementation for estimating a moving median requires tracking both the order of sample arrivals and sorting the entire buffer (window) each time a new sample is received, resulting in an algorithmic complexity of *O*(*N*^2^). A more efficient approach involves maintaining a sorted copy of the window data, allowing for the insertion of new samples into their correct positions while simultaneously deleting the oldest element in the buffer and appropriately shifting the remaining samples. With this method, originating from an ordered buffer, the complexity diminishes to *O*(*N* log *N*) operations on average. However, it demands maintaining the history of sample arrivals to the buffer, which proves both memory and time-intensive. In real-time implementations, this poses a new challenge: ensuring that the output keeps pace with the input data rate to prevent data loss.

We propose a novel real-time median estimator where the new incoming samples are inserted into the buffer in an orderly fashion. In our algorithm, instead of discarding the oldest sample, we suggest removing a sample at the extremity of the buffer that is farthest from the new sample.

Given a buffer *B* with *L* positions, being *L* = 2*m* + 1 so *x*_*m*+1_ is the element in the middle of the buffer. In Fig 1a-upper the ordered buffer is displayed with elements from 1 to 2*m* + 1 at time *k*.

**Fig 1.**
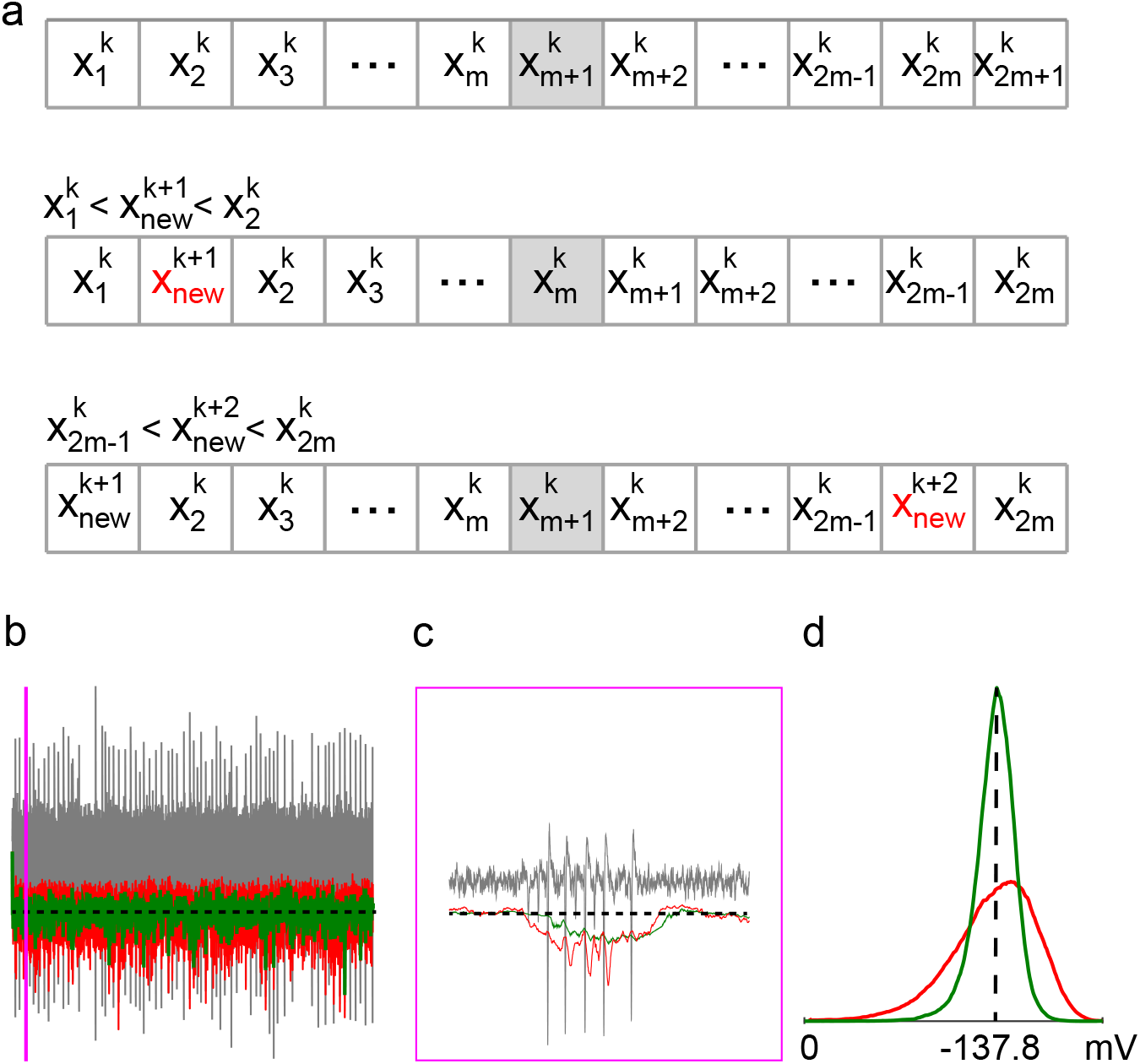
Median estimates of an extracellular recording. (a) The algorithm is illustrated during three time steps. (b) The figure shows a portion of a real extracellular recording from the prefrontal cortex of a rat [8]. The recording is displayed in gray. Thresholds estimated from the classical moving median estimator and the proposed estimator are shown in red and green, respectively. The dashed line shows the threshold computed from the whole recording as 4 *σ*_*noise*_. (c) Zoom over purple rectangle in (b). (d) The value distribution for both the classical and the new median estimators is illustrated. As can be observed, the classical median estimator has a longer tail towards negative values due to its sensitivity to outlier values resulting from spikes.

When a new sample, denoted by 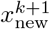, arrives at time step *k* + 1 and it is lower than the median 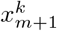, in the buffer, one of two scenarios occurs:

- Insertion within the buffer: if 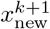 is greater than at least one element, 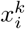 (where 1 ≤*i* ≤*m*), in the buffer, it is inserted at the appropriate position to maintain the ascending order of the buffer.
- Insertion at the beginning: if 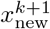 is lower than all elements in the buffer, it becomes the new minimum and is inserted at the first position.

In both cases, the current maximum element, 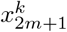, is removed from the buffer to maintain its fixed size, as showed in Fig 1a-middle. The opposite case is showed in Fig 1a-lower.

Finally, the case when the new sample is equal to the element in the center of the buffer, either the first or the last element of the buffer is dropped out randomly, and the new sample is inserted at the middle.

It is important to remark that in this algorithm: a) the elements in the buffer *B* are consistently sorted, b) the new sample *x* is always inserted in the buffer, c) There is no requirement for information about the time of sample arrival.

To determine the position where the new element should be inserted, we must either locate an element of the same value in the buffer or conduct at least *O*(log *N*) comparisons using a binary search algorithm.

## Results

While median filtering finds applications in various domains, our focus will be on its application in extracellular neuron spike detection. Electrodes in a multichannel extracellular recording setup can detect abrupt electrical changes in their vicinity.

However, they also capture the aggregate of numerous electrical signals originating from distant neuron populations, alongside various electrical artifacts.

Regardless of the individual probability distributions of these punctual sources, the central limit theorem dictates that their collective contribution approximates a normal distribution. Consequently, the stronger the electrical field perturbation caused by a nearby neuron spike on an electrode, the greater the likelihood of discerning it as a single-cell action potential. Thus, it is customary to define the signal-to-noise ratio (SNR) between spikes and background noise as the ratio of the spike peak voltage to the standard deviation of the background noise, denoted as *σ*_*noise*_. A SNR value of K indicates that the spike peak voltage is K times the standard deviation of the background noise, denoted as *σ*_*noise*_. In the case of Gaussian noise *N*(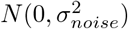), a linear relationship exists between its standard deviation and the median of the half-normal distribution *y* = |*x*|,

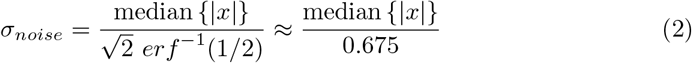

Thus, computing a threshold *T* to detect events surpassing *T* reduces the computation of *σ*_*noise*_ to the median of *y* = |*x*|, with *p*(*x*) = *N*(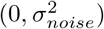).

In Fig 1, we present a schematic illustrating the step-by-step insertion of new samples into the buffer and the subsequent discarding of samples according to the algorithm outlined in the Materials and methods section. Compared to the classical moving median estimator, the proposed method exhibits a smaller variance when applied to real neuronal extracellular recordings. This reduction in variance translates to fewer missed spikes and eliminates the erroneous detection of non-existent spikes, which is particularly important for accurate spike detection thresholding.

The reduction in variance in the estimation is attributed to the fact that samples are discarded from the buffer based on their position within it, rather than their arrival times. Specifically, in a buffer initially populated with the first *L* samples in order, the classical moving median (CMM) estimator discards the oldest sample in the buffer each time a new sample arrives, whereas our algorithm (NM) discards a sample from the buffer’s extremity. Intuitively, this can be understood as a compressing process of sample selection, ultimately resulting in a buffer populated with samples closer to the median.

It is noteworthy that the classical algorithm maintains a true sampling distribution of the original signal, while ours does not fully reflect this distribution. We reasoned that while this characteristic may offer an advantage in a stationary process, it may result in a delay in reaching the new median value when a change in the median signal occurs. Furthermore, the delay in reaching the correct estimation of the median and the length of the buffer are also related. Larger buffers indeed offer less dispersion around the real median and provide more stability at the cost of slower responses to process changes. To investigate this phenomenon, we conducted a series of simulations using signals constructed from real recording noise and spikes, with spikes uniformly distributed among the basal noise. Therefore, for a set of buffers with lengths 63, 511, and 1023, we introduced a controlled change in the signal values at time 0 and examined the dynamic changes in the median estimation (see Fig 2).

**Fig 2.**
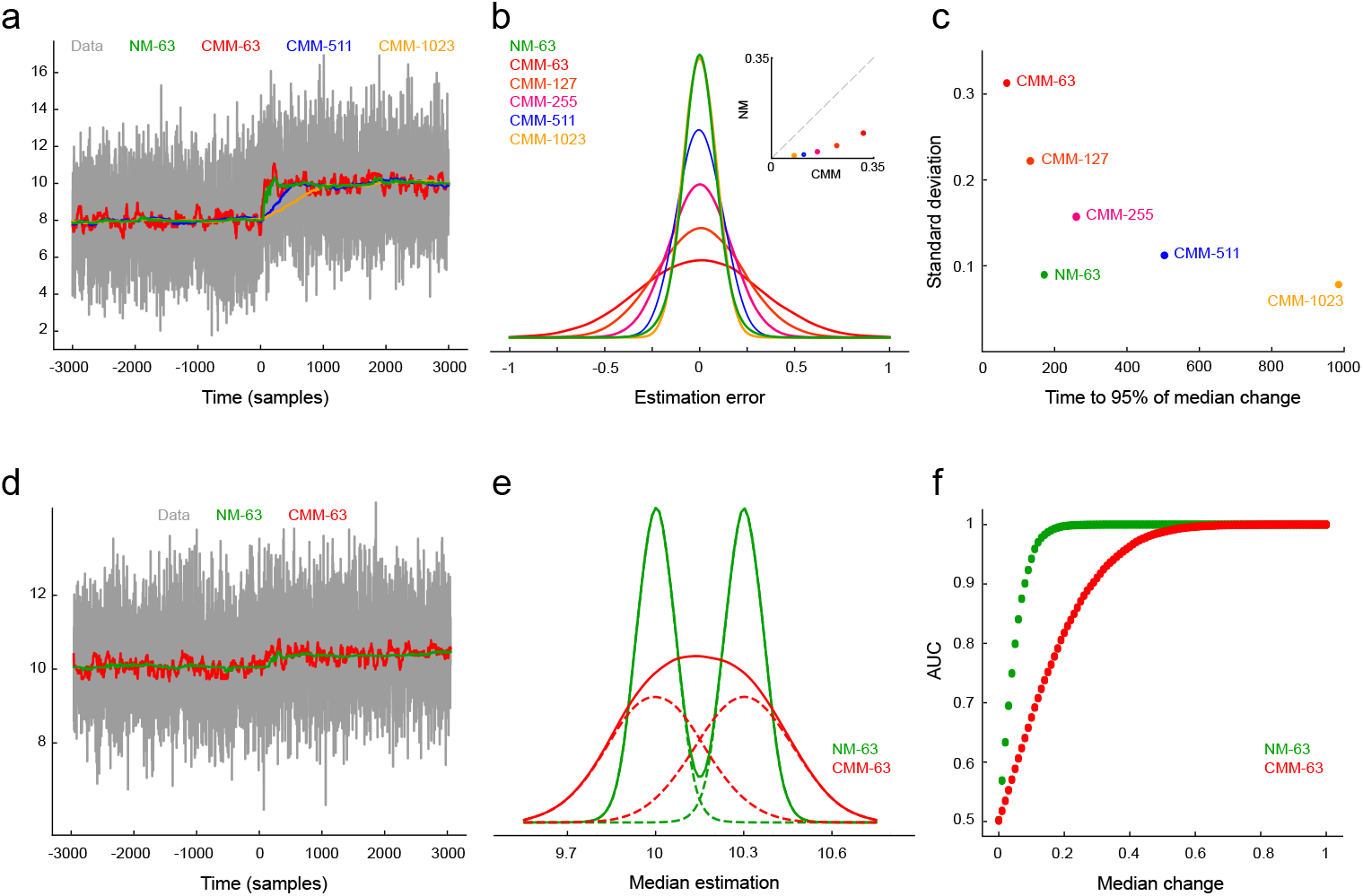
Time and spatial sensitivity to process changes. (a) A simulated extracellular recording (gray) with a median change at *t* = 0. Changes in the median estimator for buffers with lengths 63, 511, and 1023 are plotted in green, red, blue, and yellow respectively. (b) Dispersion around the true median value for different buffer lengths, including our algorithm (NM-63). (c) The compromise between dispersion and the time to reach 95% of the new median value after a change. (d),(e) Sensitivity of both algorithms using a 63-position buffer after a small change in the process. Solid lines in E depict the entire distribution of estimations before and after the change. (f) ROC analysis for changes in the median process ranging from 0 to 1.

Unlike the Classical Moving Median (CMM) estimator, where changes are reflected in the estimation within a fixed time window *L*, our algorithm’s response time depends on the magnitude of the change. For large changes with entirely new samples, our algorithm requires on average between *L/*2 and 3*L/*2 time steps to achieve the same. However, for small changes, the new samples are added to the edges of the buffer and may take longer than L steps to fully influence the estimate due to the discarding process.

Slow and gradual changes in the data dynamics may require slightly more time to converge to the true median value. While this slower reaction to changes acts as the counterpart for robustness, the method balanced this disadvantage providing a better resolution, as shown in Fig 2e.

To demonstrate the algorithm’s robustness against outliers, we analyzed the sample density dynamics within the buffer over time. To directly compare NM with the CMM algorithm, we initially filled both buffers with L pre-sorted samples. This ensures the NM estimator’s initial distribution reflects the underlying data. However, unlike CMM where this distribution remains constant, the NM estimator continuously compresses the buffer’s sample distribution towards the median value, typically within a few buffer cycles.

We ran the algorithm with a buffer length of L=1023 samples, feeding it data from a normal distribution *N*(0, 1). We then analyzed the sample density within the buffer at different points in time, starting from when the buffer was initially filled. Fig 3a shows these sample density changes. Particularly noteworthy, the initially Gaussian distribution becomes progressively more concentrated around the data’s mean (or median for a normal distribution), approaching a uniform distribution with significantly reduced variance.

**Fig 3.**
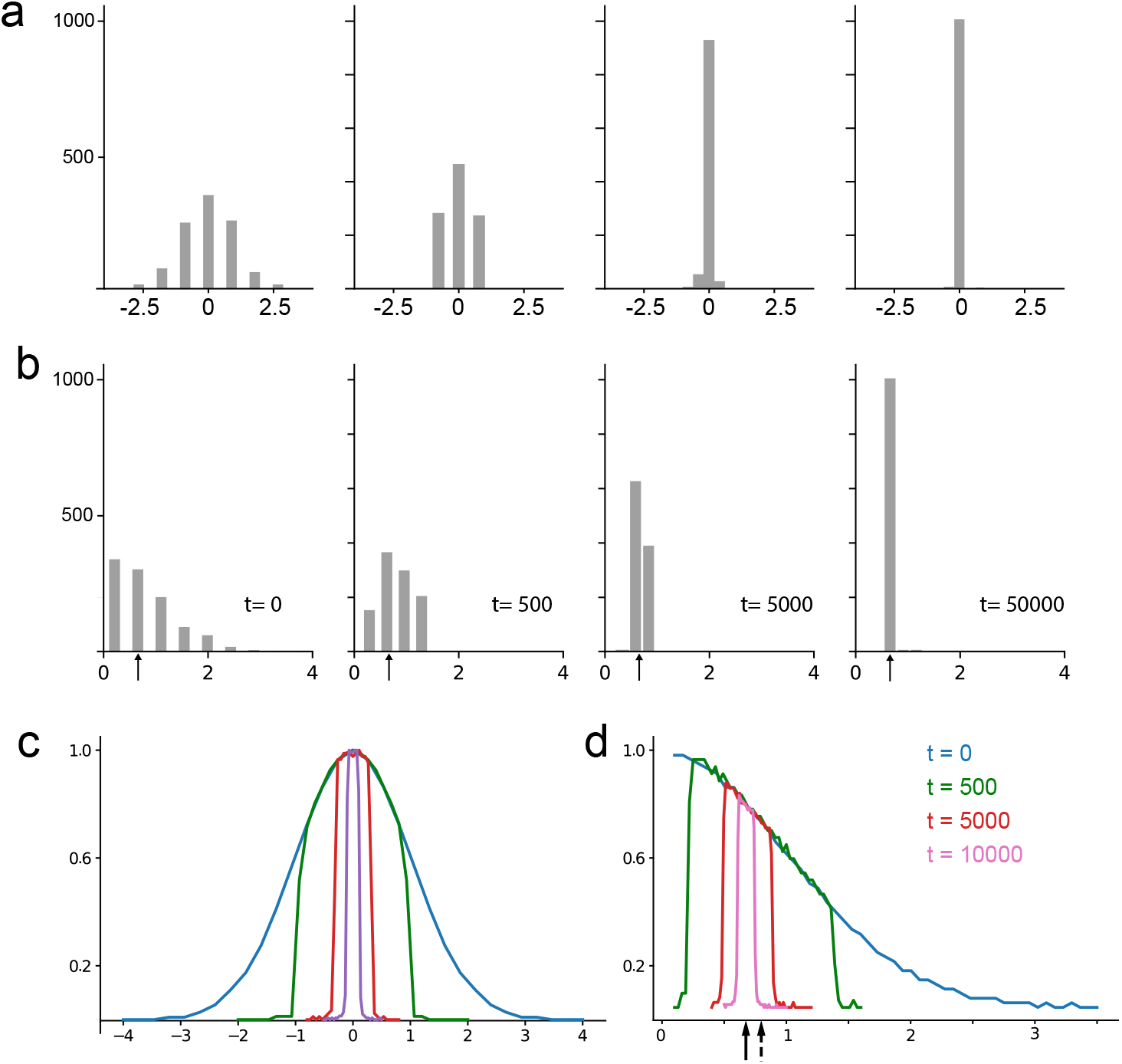
Dynamical changes in the buffer population (*L* = 1023). (a) Sample distribution across time for *x∼ N*(0, 1). (b) Sample distribution across time for |*x*|, when *x ∼ N*(0, 1), the arrow shows the true signal median. (c),(d) The sample distribution of the data across the buffer and over time is visualized (10,000 repetitions average) of the experiment for *x* and |*x*|, when *x ∼ N*(0, 1). Results were normalized in order to observe the magnification effect around the true signal’s median value as time progresses. Continuous arrow: true signal median. Dashed arrow: true signal mean

We repeated the analysis for a folded normal distribution (the distribution of |*x* |when *x∼ N*(0, 1). Fig 3b shows the sample distribution within the buffer over time, starting from a buffer completely filled with fresh samples drawn from the folded normal distribution, and continuing up to 50, 000 time steps. This scenario, with a folded normal distribution, is more relevant to real extracellular recordings where the median and mean typically differ. As can be readily observed, the buffer samples distribute around the noise median in this case.

Interestingly, repeating both experiments 10,000 times and averaging their sample distributions across different time points reveals a consistent trend. The average sample distribution concentrates around the median value of the initial distribution, essentially copying its shape. This behavior is evident in Fig 3c for the normal distribution and Fig 3d for the folded normal distribution. Consequently, filling the buffer with values increasingly closer to the true median with each iteration enhances the method’s robustness against outlier perturbations. As Fig 3 illustrates, the buffer prioritizes samples closer to the median value as time goes. This contrasts with the classical moving median estimator, where an outlier resides in the buffer for *L* steps. In our method, outliers are relegated to the buffer’s extremes and have a significantly lower probability of remaining influential for many steps, effectively halving with each step k, 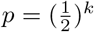.

By definition, the median *m* is the value that satisfy

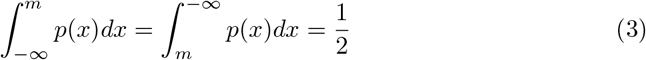

for any other value *m*_*x*_ different to *m* we will have

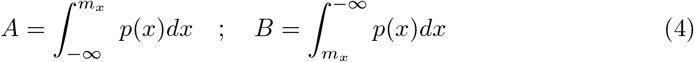

with *A* ≠ *B*.

If the value at the center of the buffer *m*_*x*_ is less than the true median *m*, there will be an imbalance. The distribution function, *F*(*x*), tells us that there are more samples greater than *m*_*x*_ than expected *F*(*m*) *− F*(*m*_*x*_) *>* 0. These additional high-value samples on the right side of the buffer tend to push *m*_*x*_ towards the left. This creates an opportunity for values greater than *m*_*x*_ to occupy the buffer center position. The opposite happens when *m*_*x*_ is larger than the true median.

This process can be likened to the diffusion of non-charged particles across a membrane, as described by Fick’s Laws [9]:

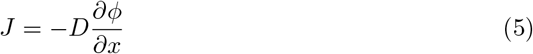

where *J* is the net flux across the membrane, *D* is the diffusion coefficient (*D* = 1 here) and 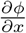 is the concentration difference across the membrane.

The process reaches equilibrium, indicated by a zero net flux of zero particles across the membrane, when the concentration of particles on both sides becomes equal. In simpler terms, this occurs when the same proportion of samples are found above and below the central value, *A* = *B* = 1*/*2, which occurs when *m*_*x*_ itself equals the true median *m*.

By exploiting this natural dynamic, our algorithm achieves an unbiased estimation of the signal’s median under stationary conditions.

## Discussion

There are two main categories of quantile estimators: those that calculate the statistics using the entire dataset and those that rely on a subset of the data.

There is a significant amount of research on computing median estimators in data streams, specifically to be applied to filtering techniques. These techniques often involve calculating the exact median of a window containing L samples and then using this value to estimate an appropriate threshold.

On the other hand, quantile estimation is a powerful technique for characterizing dataset properties. It enables the online estimation of various order statistics, including the minimum, maximum, median, quartiles, and any other quantiles (q-quantile).

In streaming algorithms, finding the minimum and maximum values only requires *O*(1) operations, while estimating the median requires at least *O*(log *N*) operations [10, 11]. Efficient online computation of quantile statistics are mainly based on the work of Greenwald and Khanna [12, 13]. These algorithms are designed to maintain statistics of arbitrary quantiles. While they are efficient in terms of memory usage and have optimal complexity performing order *O*(1) operations, these operations are not suitable for implementation on field-programmable logic arrays (FPGAs) or Application-Specific Integrated Circuit (ASICs). FPGAs are often preferred for their low-power and high-performance capabilities in real-time applications, making them unsuitable for these specific algorithms. Consequently, the complexity and computational cost of implementing them outside of a traditional computer or embedded system are very high.

Although the proposed method is limited to estimating the median, its implementation on FPGAs for this specific task is remarkably straightforward. This efficiency stems from the fact that it only requires basic operations as comparisons and shifting, which are highly optimized for FPGAs and GPUs. Furthermore, as parallelization is the main strength of FPGA/GPU, comparison with all elements of the buffer can be done in O(1) [2] and shifting elements in the buffer is also O(1), which reduces the system clock to twice the sampling frequency (instead of *O*(log *N*)), thus reducing power consumption.

## Conclusion

We presented a novel method to estimate the median of a data stream. Our method relays on a buffer of arbitrary length. For a given length our algorithm will always provide a higher accuracy estimation than a classical moving median estimator while avoiding to store the age of each sample in the buffer

